# Collecting Duct-Targeted Lipid Nanoparticles Deliver *Pkd2* mRNA to Restore Polycystin-2 and Attenuate ADPKD

**DOI:** 10.64898/2026.06.02.728835

**Authors:** Joshua Giblin, Karla Lambaren, Ronnie LaMastro, Isabella Suzuki, Yi Huang, Rowan Simon, Jose Zarate-Diaz, Ali Osouli, Jessica Pham, Kenneth Hallows, Eun Ji Chung

## Abstract

Autosomal dominant polycystic kidney disease (ADPKD), a leading genetic cause of kidney failure, is caused by mutations in the *PKD1* or *PKD2* genes, resulting in functional polycystin 1 (PC1) or polycystin 2 (PC2) deficiency, and is characterized by progressive cyst expansion in the kidneys. However, no approved therapy directly restores polycystin expression, and current treatments target downstream pathways rather than the genetic defect. Gene replacement therapy offers a direct route to functional rescue, but efficient delivery to cyst-lining renal epithelia remains a major barrier. To meet this challenge, we developed a lipid nanoparticle (LNP) that incorporates a collecting duct (CD) targeting peptide (CD LNPs) to enable delivery of *Pkd2* mRNA (CD-m*Pkd2*) to renal CD epithelia, the predominant site of cyst origin. CD LNPs increased renal accumulation and targeted CD cells *in vivo*, outperforming non-targeted formulations. In a *Pkd2*-deficient mouse model, repeated administration of CD-m*Pkd2* induced reversal of established cystic disease, restored renal architecture, and reduced the fibrotic and inflammatory microenvironment characteristic of ADPKD. Furthermore, CD-m*Pkd2* was well tolerated without detectable off-target toxicity. Given that PC2 supplementation can attenuate disease in *Pkd1*-deficient models, we further demonstrate that a single dose of CD-mPkd2 reduces cyst burden and improves renal function across this distinct genetic background. These findings establish CD-m*Pkd2* as a potential therapeutic strategy for ADPKD across genetic backgrounds.

## Introduction

Autosomal dominant polycystic kidney disease (ADPKD) is the most common inherited kidney disorder, affecting more than 12 million individuals worldwide^1^. The disease is characterized by progressive cyst formation, renal enlargement, and eventually kidney failure, with patients reaching end-stage renal disease (ESRD) by 50 years of age^1,2^. Beyond kidney failure, patients frequently develop urinary infections, kidney stones, hypertension, and chronic and acute abdominal pain^1,3^.

ADPKD arises from mutations in either the *PKD1* (∼80% of cases) or *PKD2* (∼15% of cases) genes, encoding polycystin-1 (PC1) and polycystin-2 (PC2), respectively^1,4^. These proteins form the polycystin complex that regulates intracellular calcium signaling, cellular metabolism, and tubular renal epithelial architecture^5,6^. Disruption of functional polycystins alters intracellular calcium homeostasis and promotes cyclic adenosine monophosphate (cAMP)–dependent proliferation and fluid secretion in renal cells, driving cyst expansion^7–9^. Although cysts arise from all nephron segments, the collecting duct (CD) epithelium represents a predominant site of cyst initiation and expansion, making it an attractive cellular target for therapeutic intervention^10–13^.

Currently, the only FDA approved therapy is tolvaptan, a vasopressin V2 receptor (V2R) antagonist that inhibits V2R-mediated cAMP signaling in CD epithelia^14^. By lowering intracellular cAMP, tolvaptan reduces cyst epithelial proliferation and fluid secretion, thereby slowing total kidney volume growth and modestly delaying decline in renal function^14,15^. However, its use is limited by aquaretic side effects, hepatotoxicity risk, and high discontinuation rates^16–18^. Importantly, tolvaptan does not address the underlying polycystin deficiency and the genetic root of ADPKD. Thus, there remains a critical need for therapies that directly correct or compensate for the loss of polycystin signaling.

Gene replacement therapy represents a strategy for monogenic diseases such as ADPKD. Genetic re-expression of *Pkd2* using an inducible allele system in which gene expression was restored after disease onset has been shown to reverse cystic kidney disease in mice, demonstrating that restoration of polycystin function can reverse established pathology^19^. Furthermore, in ADPKD, *PKD1* and *PKD2* translation is suppressed by microRNA-17 (miR-17) and previous work has demonstrated that inhibiting miR-17-mediated repression on the *Pkd2* transcript increases PC2 expression and attenuates cyst progression in *Pkd1*-deficient models, highlighting that PC2 replacement or supplementation may represent a broadly applicable approach across genetic subtypes of ADPKD^20,21^. Messenger RNA (mRNA) therapeutics delivered by lipid nanoparticles (LNPs) offer a clinically validated platform for transient protein replacement^22^. However, efficient mRNA delivery to extrahepatic organs such as the kidney remains a major translational barrier^23^. Achieving therapeutically meaningful mRNA delivery to cyst-lining renal epithelial cells has not been demonstrated to date.

Here, we engineered CD–targeted lipid nanoparticles (CD LNPs) functionalized with a peptide ligand to enhance uptake by cyst-forming CD epithelial cells. We hypothesized that delivery of *Pkd2* mRNA to these cells would restore PC2 expression, decrease cyst-associated signaling, and attenuate or reverse cyst growth. Using inducible models of *Pkd2*-deficient ADPKD, we evaluated renal biodistribution, cell-type specificity, therapeutic efficacy, and systemic safety following repeated systemic administration. CD-targeted LNPs enhanced renal accumulation and targeted CD epithelia *in vivo*, enabling restoration of PC2 expression and reversal of established cystic disease in *Pkd2*-deficient mice. Building on prior work demonstrating that PC2 supplementation attenuates disease in *Pkd1*-deficient models, we evaluated CD-m*Pkd2* in an inducible *Pkd1*-deficient model of ADPKD^20^. In this model, a single dose of CD-m*Pkd2* attenuated cyst progression and improved renal function, although full normalization to healthy controls was not achieved. Together, these findings establish a mRNA replacement strategy for the first time and support the feasibility of polycystin restoration or supplementation therapy broadly across ADPKD genetic backgrounds using CD-targeted LNPs.

## Results

### Identification of kidney cell transfecting LNP formulations and optimization for collecting duct targeting

The relationship between LNP lipid composition and renal cellular tropism remains poorly defined, as systemic LNP platforms exhibit dominant hepatic accumulation, limiting rational design of kidney-targeted formulations for mRNA delivery^24–26^. Guided by prior observations that DOTAP-containing LNPs exhibit enhanced renal accumulation^24,27^, we constructed a focused library of four formulations altering the molar ratios of DOTAP (20-35 mol%) and DSPE-PEG(2000)-methoxy (1.5-5 mol%). Furthermore, to expand our library, LNPs were synthesized with either D-lin-MC3-DMA (MC3) or ALC-0315 (ALC), ionizable lipids used in clinically approved RNA therapeutics. All LNPs were formulated with mCherry mRNA at a 10:1 ionizable lipid–to–mRNA weight ratio and characterized for encapsulation efficiency, hydrodynamic diameter, zeta potential, and polydispersity index (PDI). All formulations, except ALC-4, achieved > 80% encapsulation (**Fig. 1A, B**). Particle diameters ranged from 84–117 nm (**Fig. 1C**), zeta potentials from −5 to +11 mV depending on DOTAP content (**Fig. 1D**), and PDI values were < 0.3 (**Fig. 1E**).

**Figure 1.**
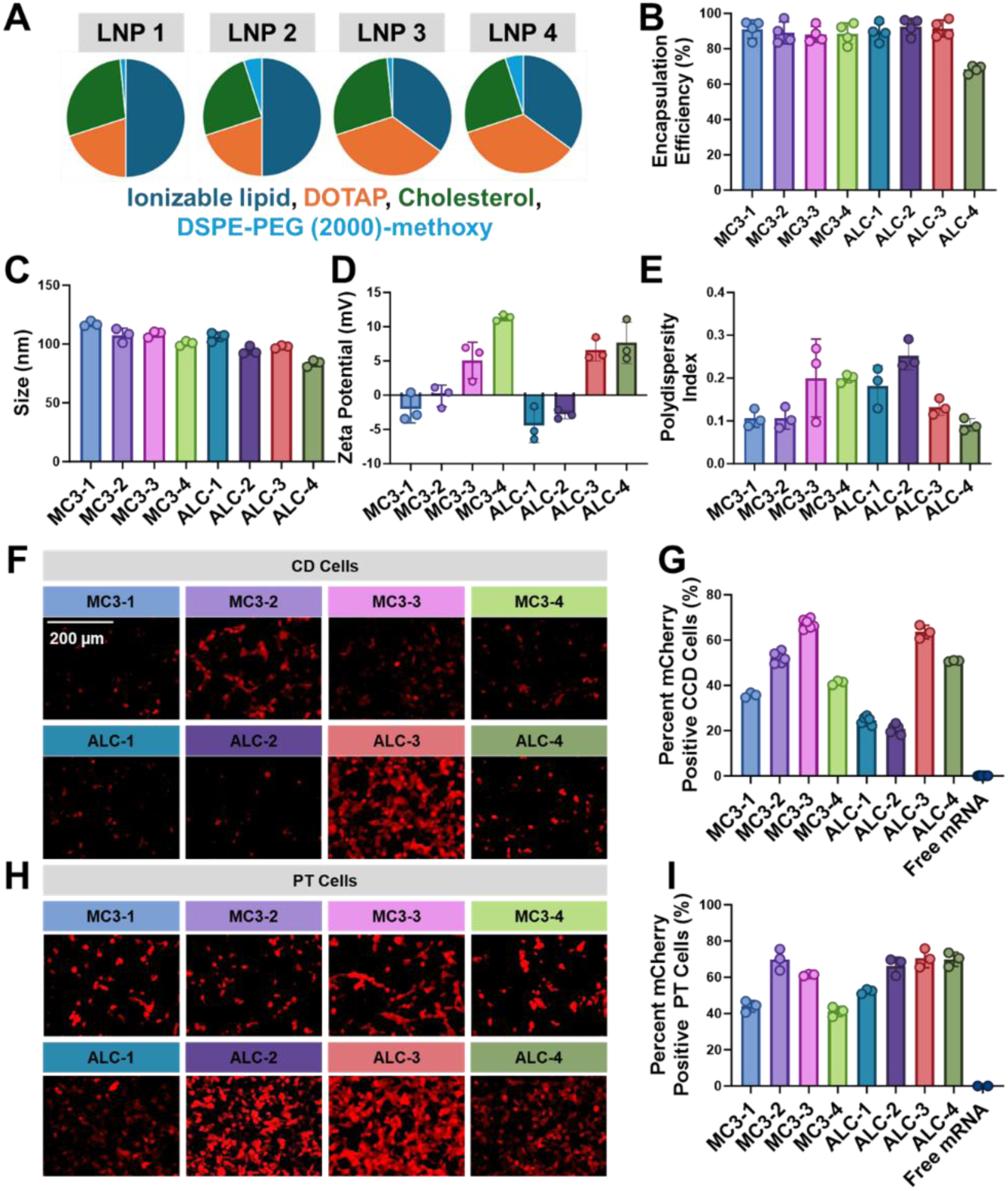
Synthesis and screening of LNP formulations for mRNA delivery to renal tubular epithelial cells. (A) Pie charts depict the four formulations that were synthesized based on varying DOTAP and DSPE-PEG(2000)-methoxy content. (B) mRNA encapsulation efficiency, (C) size, (D) zeta potential, and (E) polydispersity index values across the eight LNPs. (F) Representative fluorescence images of CD cells following transfection with mCherry-encoding mRNA delivered by MC3- and ALC-based LNP formulations. Scale bar = 200 μm. (G) Quantification of mCherry-positive CD cells following LNP treatment (500 ng/mL mCherry mRNA; 24 hours). (H) Representative fluorescence images of proximal tubular epithelial (PT) cells following transfection with mCherry-encoding mRNA delivered by MC3- and ALC-based LNP formulations. (I) Quantification of mCherry-positive PT cells following LNP treatment (500 ng/mL mCherry mRNA; 24 hours). Data are represented as mean ± SD. Statistical analysis was calculated with a one-way ANOVA with Tukey’s post hoc test.

Transfection efficiency was evaluated in mouse proximal tubular (PT) epithelial cells and mouse CD cells, the principal epithelial populations contributing to cyst formation in ADPKD^10,28^. Cells were incubated with LNPs (500 ng/mL mCherry mRNA) for 24 hours. In CD cells, MC3-3 and ALC-3 (35 mol% ionizable lipid, 35 mol% DOTAP, 28.5 mol% cholesterol, and 1.5 mol% DSPE-PEG(2000)-methoxy) produced the highest transfection (**Fig. 1F, G**). In PT cells, transfection was generally higher across formulations, with ALC-3 producing the greatest transfection efficiency (**Fig. 1H, I**). Thus, the ALC-3 formulation was selected as the base formulation and subsequently functionalized with a CD-targeting peptide to enhance specificity for cyst-forming CD epithelia.

To direct LNPs to cyst-lining CD cells, the primary site of cystogenesis in ADPKD, we functionalized LNPs with a CD-targeting peptide by substituting DSPE-PEG(2000)-methoxy with the corresponding DSPE-PEG(2000) CD-targeting peptide conjugate (**Fig. 2A**)^10,29^. LNPs formulated with 1.5 mol% DSPE-PEG(2000)-CD targeting peptide, exhibited the highest transfection in CD cells, without altering transfection in PT cells relative to non-targeted (NT) LNPs (**Fig. 2B-D**). Importantly, incorporation of the CD-targeting peptide onto the LNPs (hereafter CD LNPs) did not affect cellular metabolic activity at doses up to 1000 ng/mL mRNA, indicating minimal cytotoxicity across and beyond the functional dosing range used in this study. Specificity of CD-targeting peptide–mediated LNP uptake was confirmed by competitive inhibition, as pre-incubation with free peptide reduced uptake of DiR-labeled CD LNPs by 2.1-fold compared to untreated controls, an effect not observed with NT LNPs, confirming that uptake is mediated specifically through the CD-targeting peptide (**Fig. 2E**). Consistent with enhanced targeting, CD LNPs mediated significantly (p < 0.05) higher mCherry expression than NT LNPs across multiple doses in inner medullary collecting duct (IMCD) cells (**Fig. 2F, G**), with enhanced expression evident as early as 4 hours and peaking at 48 hours (**Fig. 2H, I**).

**Figure 2.**
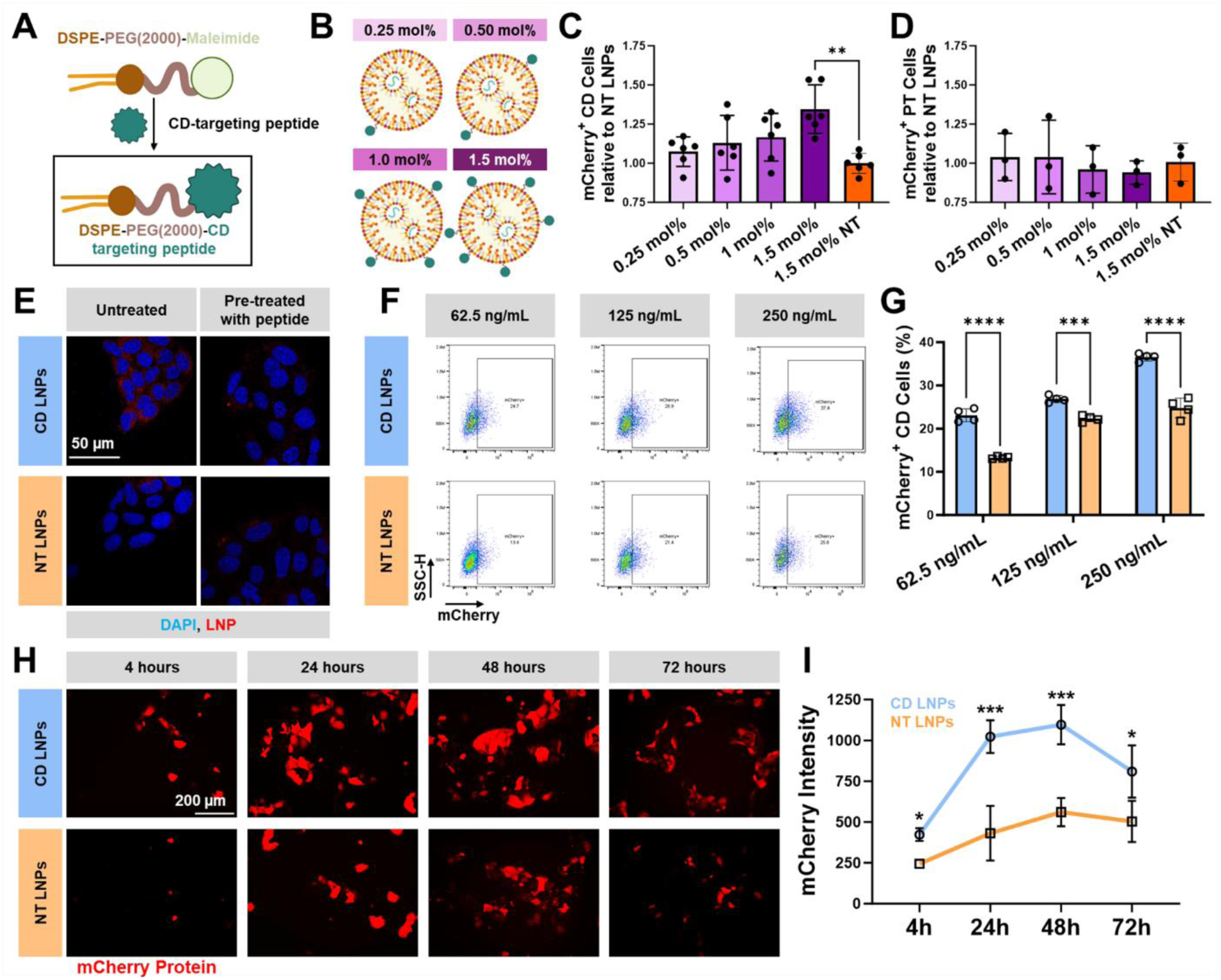
LNPs functionalized with a CD-targeting peptide achieve enhanced CD-targeting *in vitro*. **(A)** Schematic representation of the conjugation between DSPE-PEG(2000)-maleimide, the collecting duct (CD)-targeting peptide, and the resulting DSPE-PEG(2000)-CD targeting peptide. **(B)** Illustrations of LNPs containing various mol% of DSPE-PEG(2000)-CD targeting peptide. **(C)** Quantification of mCherry-positive CD and **(D)** PT cells after transfection and normalized to mCherry-positive cells after NT LNP treatment (24 hours; 250 ng/mL mCherry mRNA). **(E)** Fluorescence images of CD cells incubated with DiR-labeled CD or NT LNPs following pre-treatment with the CD-targeting peptide. Scale bar = 50 µm. **(F)** Flow cytometry and **(G)** quantification of mCherry-positive cells after LNP treatment with various doses after 24 hours. **(H)** Fluorescence images of CD cells following treatment with CD or NT LNPs from 4-72 hours. Scale bar = 200 µm. **(I)** Quantification of mCherry mRNA delivery at each timepoint. Data are represented as mean ± SD. Statistical analysis was calculated with a student’s *t* test. ** p < 0.01, *** p < 0.001, **** p < 0.0001.

### Efficient kidney accumulation and CD-targeting by CD LNPs

To evaluate the *in vivo* biodistribution and renal targeting efficiency of CD LNPs in a disease-relevant setting, we utilized the *Pkd2*^fl/fl^; Pax8-Cre inducible mouse model of cystic kidney disease (**Fig. 3A**)^30^. Following doxycycline-mediated induction from postnatal day (P) 28-31, mice received a single intravenous injection of DiR-labeled CD LNPs or NT LNPs at P125. *Ex vivo* imaging of isolated kidneys, 24 hours post-administration, revealed significantly higher accumulation of CD LNPs compared with NT LNPs (**Fig. 3B**). Quantification of DiR fluorescence intensity confirmed a 36.1% increase in kidney signal in CD LNP-treated mice compared to NT LNP controls (p < 0.001) (**Fig. 3C**). Moreover, organ-level biodistribution analysis revealed a 1.8-fold increase in kidney enrichment for CD LNPs relative to NT LNPs (**Fig. 3D**).

**Figure 3:**
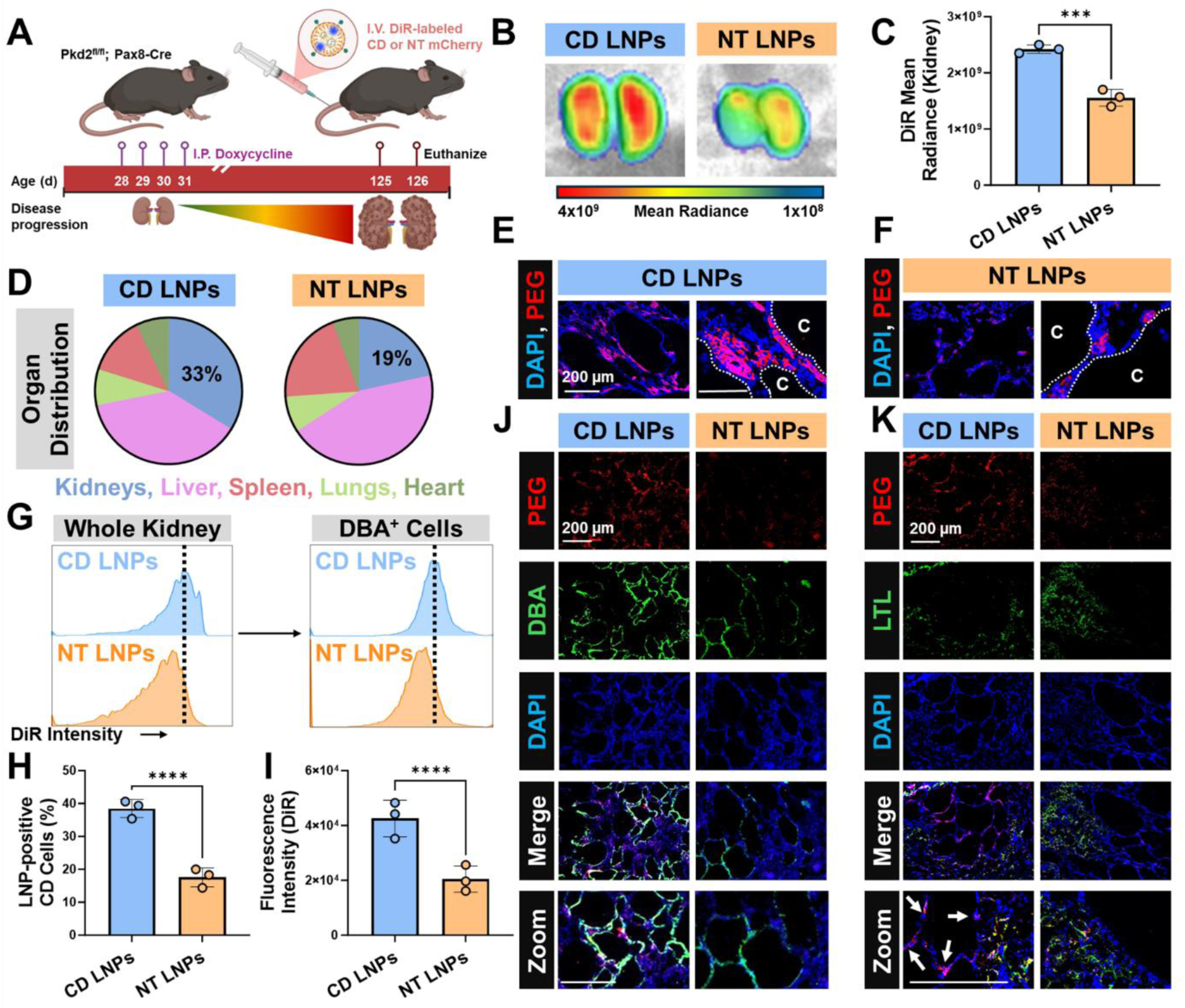
CD-targeting LNPs accumulate in renal collecting duct cells in mice with cystic kidney disease. **(A)** Experimental schematic. *Pkd2*^fl/fl^; Pax8-Cre mice were treated with doxycycline to induce *Pkd2* inactivation and cyst formation. DiR-labeled LNPs encapsulating mCherry mRNA were administered intravenously (I.V.), and mice were euthanized for tissue analysis. **(B)** Representative *ex vivo* fluorescence images of kidneys harvested 24 hours following treatment with CD or NT LNPs. **(C)** Quantification of DiR fluorescence intensity in kidneys following LNP administration (N = 3). **(D)** Distribution of DiR fluorescence across major organs (kidneys, liver, spleen, lungs, heart) following systemic administration of LNPs. **(E-F)** Fluorescence images of kidney sections stained for PEG to detect LNPs (red) and DAPI (nuclei). Scale bar = 200 µm. **(G)** Histograms of DiR intensity in whole kidney digest and *Dolichos biflorus* agglutinin (DBA)-positive cells. **(H)** Quantification of the percentage of DiR (LNP)-positive DBA-positive (CD) cells in dissociated kidneys. **(I)** Quantification of the DiR fluorescence intensity in DBA-positive cells in dissociated kidneys. **(J)** Immunofluorescence images of kidney sections stained for PEG (red), a CD marker (DBA, green), and DAPI (nuclei). **(K)** Immunofluorescence images of kidney sections stained for PEG (red), a proximal tubular epithelial cell marker (LTL, green), and DAPI (nuclei). Scale bar = 200 µm. Data are represented as mean ± SD. Statistical analysis was calculated with a student’s *t* test. *** p < 0.001, **** p < 0.0001.

To examine intrarenal LNP distribution, kidney sections were stained for PEG and analyzed by fluorescence microscopy. Strong LNP signal was detected along the cyst epithelium in mice treated with CD LNPs, whereas kidneys from NT LNP–treated mice exhibited weaker and more diffuse signal, mainly localized to pericystic regions rather than cyst-lining epithelial cells (**Fig. 3E–F**). Because CD cells are a major site of cyst formation, we next assessed LNP uptake in DBA-positive CD cells. Flow cytometric analysis of dissociated kidneys revealed there was a 2.2-fold higher percentage of LNP-positive cells within the DBA⁺ epithelial population in CD LNP–treated mice compared with NT LNP controls (**Fig. 3G, H**). Consistent with this, CD LNP treatment resulted in a 2.1-fold higher DiR fluorescence intensity in DBA-positive CD cells (**Fig. 3G, I**). To further assess intrarenal localization, kidney sections were co-stained with markers for CD cells (DBA), PT cells (LTL), and glomeruli (WGA lectin)^31^. Imaging showed substantial co-localization of CD LNP signal with CD cells, whereas NT LNP signal showed limited overlap with CD cells (**Fig. 3J**). PT cells exhibited comparable LNP uptake across both treatment groups, indicating similar access to this compartment (**Fig. 3K**). In kidney sections stained with WGA lectin to identify glomerular structures, neither CD LNPs nor NT LNPs were found (**Fig. S1A**). Together, these findings demonstrate that CD LNPs enhance renal accumulation and target CD epithelia, including cyst-lining cells, and that the expanded cystic CD compartment in ADPKD amplifies the proportion of disease-relevant cells reached, a key advantage for targeted therapeutic delivery.

### Repeated CD-mPkd2 administration reverses established renal cystic disease in Pkd2-deficient mice

The preferential accumulation of CD LNPs within cyst-lining epithelia suggested that this platform could serve as a vehicle for targeted mRNA delivery in ADPKD. We therefore loaded CD LNPs with full-length mouse *Pkd2* mRNA (m*Pkd2*) containing a 300-adenosine poly(A) tail. We found through a gel shift assay that a 20:1 ionizable lipid–to–mRNA weight ratio prevented the mRNA from migrating down the gel (**Fig. S2A**), indicating successful entrapment of full-length *Pkd2* mRNA. CD-m*Pkd2* nanoparticles measured 106.5 nm in diameter, with +7.8 mV zeta potential, PDI of 0.16, 84% encapsulation efficiency, and a spherical morphology with no reduction in cell viability (**Fig. S2B-F**).

To determine whether CD-m*Pkd2* could reverse established renal cystic disease, *Pkd2*^fl/fl^;Pax8-Cre mice were induced at P28–31 and allowed to develop cystic disease until P99. Beginning at P99, a timepoint at which kidneys exhibit substantial cyst burden, mice received CD-m*Pkd2* every three days at 1 mg/kg of *Pkd2* mRNA or PBS control (**Fig. 4A**). At endpoint (P128), gross examination revealed that CD-m*Pkd2* treatment reduced the kidney weight-to-body weight (KW/BW) ratio by 62% relative to PBS-treated mice (**Fig. 4B, C**). Notably, kidneys from the CD-m*Pkd2*-treated cohort were 31% smaller than at baseline (P99), indicating true regression of established disease rather than attenuation of progression (**Fig. 4B, C**).

**Figure 4:**
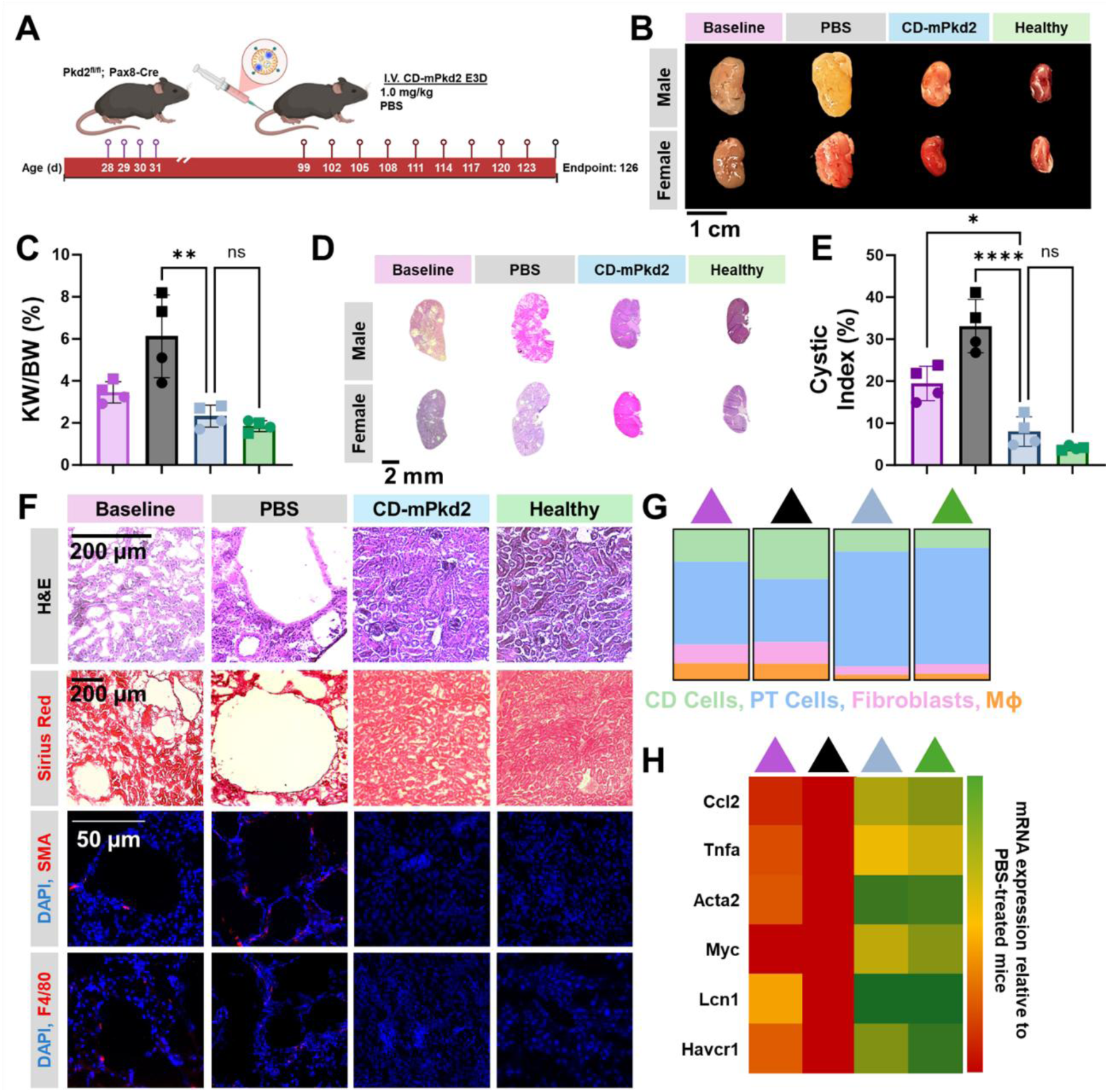
CD-m*Pkd2* treatment reverses cyst burden and improves kidney pathology in a *Pkd2*-deficient mouse model. **(A)** Experimental schematic. *Pkd2*^fl/fl^; Pax8-Cre mice were induced with doxycycline from P28-31. Mice received intravenous (I.V.) injections of CD-m*Pkd2* at 1.0 mg/kg or PBS beginning at P99 every three days until P128. **(B)** Gross kidney images from male and female mice at baseline (P99) and after treatment (P128). Uninduced age-matched mice are shown for comparison. Scale bar = 1 cm. **(C)** Quantification of KW/BW across treatment groups. **(D)** Whole kidney sections from male and female mice after different treatments. Scale bar = 2 mm. **(E)** Quantification of cystic index in the kidney after various treatments. **(F)** Representative kidney histology and immunostaining across treatment conditions. Top row, hematoxylin and eosin (H&E) staining. Second row, Sirius Red staining indicating collagen deposition. Third row, immunofluorescence staining for α-smooth muscle actin (SMA; red) with DAPI (blue) to visualize activated myofibroblasts. Bottom row, immunofluorescence staining for F4/80 (red) with DAPI (blue) to detect macrophage infiltration. Scale bars = 50 μm. **(G)** Kidney cell composition in the indicated groups determined by flow cytometry. (n = 3) **(H)** Heatmap of RT–qPCR analysis showing expression of inflammatory and injury-associated genes (*Ccl2, Tnfa, Acta2, Myc, Lcn1, Havcr1*) in kidneys from treated mice relative to PBS controls (n = 3). Data are represented as mean ± SD. Statistical analysis was calculated with a one-way ANOVA with Tukey’s post hoc test. ns = not statistically significant, * p < 0.05, ** p < 0.01, **** p < 0.0001.

In addition to kidney enlargement, histological analysis of kidneys from mice treated with CD-m*Pkd2* demonstrated a significant improvement in renal architecture compared to kidneys from PBS-treated mice. Specifically, CD-m*Pkd2* treatment reduced cystic index by 76% compared to PBS-treated mice and 58% relative to baseline, indicating reversal of established disease (**Fig. 4D, E**). Notably, kidneys from the CD-m*Pkd2* group exhibited compact tubular organization and minimal tubular dilation, closely resembling healthy controls (**Fig. 4F**).

Immunofluorescence staining of kidney tissue revealed strong PC2 signal in tubular epithelial cells of healthy kidneys, whereas PBS-treated cystic kidneys exhibited markedly reduced PC2 expression (**Fig. S3**). Co-staining with nephron segment markers revealed restoration of PC2 in CD-m*Pkd2*-treated mice in DBA-positive CD cells and LTL-positive PT cells. In CD-m*Pkd2*-treated mice, PC2 signal was broadly distributed along tubular epithelia and closely resembled the pattern and intensity observed in healthy kidneys (**Fig. S3**). These findings indicate that repeated CD-m*Pkd2* administration restored PC2 expression across multiple nephron segments.

Fibrotic and inflammatory remodeling was also reduced following treatment. Sirius Red staining revealed decreased pericystic collagen deposition in both treatment groups, with levels in the CD-m*Pkd2* group comparable to healthy kidneys (**Fig. 4F**). Furthermore, immunofluorescence revealed smooth muscle actin (SMA)–positive myofibroblasts and F4/80-positive macrophage infiltration, with the CD-m*Pkd2*-treated kidneys exhibiting inflammatory and fibrotic profiles similar to healthy tissue (**Fig. 4F**).

To track disease progression and treatment-induced kidney remodeling across timepoints, we compared renal cellular composition in healthy controls, *Pkd2*-deficient mice at treatment initiation (P99), and endpoint PBS- or CD-m*Pkd2*-treated mice (P128) (**Fig. 4G**). Compared to uninduced healthy controls, kidneys from *Pkd2-*deficient mice at baseline (P99) and *Pkd2-*deficient PBS-treated mice at endpoint (P128) exhibited a 2.3-fold and 4.5-fold increase in CD cells (6.2% vs.14.1% vs. 27.8%), a 2.5-fold and 3.7-fold increase in fibroblasts (3.2% vs. 8.1% vs. 11.9%), a 3.4-fold and 4.4-fold increase in macrophages (2.1% vs. 7.2% vs. 9.3%), along with a 14% and 17% decrease in PT cells (41.1% vs. 35.7% vs. 34.2%) (**Fig. 4G**). CD-m*Pkd2* treatment reduced the proportions of CD cells (8.2%), fibroblasts (3.1%), and macrophages (1.9%), while increasing PT cells (42.0%) to levels observed in healthy kidney tissue (p > 0.05 for all comparisons) (**Fig. 4G**). These findings are consistent with the remodeling of the renal microenvironment following CD-m*Pkd2* treatment, characterized by reduced fibrosis and inflammation and recovery of epithelial cell composition. Molecular analysis using RT-qPCR of kidney lysate paralleled these findings. In CD-*mPkd2*-treated mice there were significant reductions in inflammatory, cyst-associated, and injury-related transcripts, including *Ccl2*, *Tnfa*, *Acta2*, *Myc*, *Lcn1*, and *Havcr1* compared to PBS-treated mice at P128 (**Fig. 4H**). Notably, in the kidneys of CD-m*Pkd2*-treated mice, expression of these genes approached levels observed in healthy kidneys, indicating attenuation of inflammatory, fibrotic, and cystogenic gene programs (**Fig. 4H**). Together, these data demonstrate that repeated CD-m*Pkd2* administration reprograms the cystic renal microenvironment, restoring structural, cellular, and molecular features characteristic of healthy kidney tissue.

Importantly, repeated CD-m*Pkd2* administration was well tolerated. Serum aspartate aminotransferase (AST) and alanine aminotransferase (ALT) levels were comparable to healthy controls and not elevated relative to PBS-treated mice (**Fig. S4A, B**). Histological examination of liver, lung, spleen, and heart revealed no evidence of architectural disruption or overt tissue injury (**Fig. S4C**). These findings indicate that repeated systemic administration of CD-m*Pkd2* at therapeutically effective doses does not induce detectable off-target toxicity.

### Supplementation of PC2 through CD-mPkd2 slows cyst growth in a preclinical Pkd1-deficient mouse model of ADPKD

Given prior evidence that increasing PC2 levels ameliorates disease features in *Pkd1*-deficient models, we evaluated whether CD-m*Pkd2* could mitigate *Pkd1*-driven cystogenesis^20^. We first assessed CD-m*Pkd2* in 3D IMCD *Pkd1* knockout cyst cultures. Treatment with CD-m*Pkd2* (250 ng/mL) significantly reduced cyst diameter by 25% at 48 hours and 35% at 96 hours relative to PBS controls. Neither empty LNPs nor free m*Pkd2* produced comparable effects, indicating that nanoparticle-mediated delivery was required for functional rescue (**Fig. 5A-C**).

**Figure 5:**
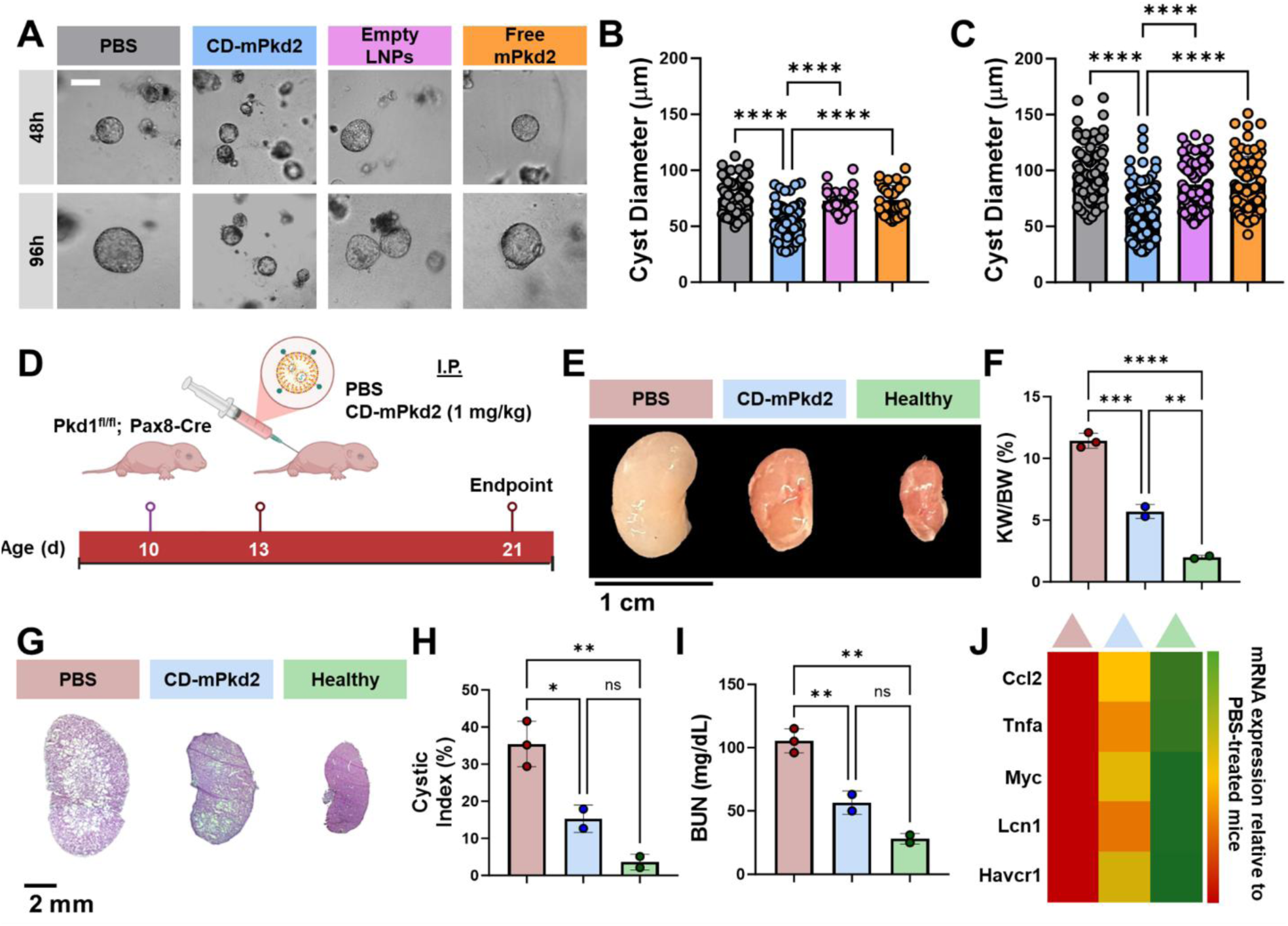
CD-m*Pkd2* suppresses cyst growth *in vitro* and improves kidney function in a *Pkd1*-deficient mouse model. **(A)** Brightfield images of *Pkd1*-knockout IMCD cells at 48 hours and 96 hours after treatment. Scale bar = 100 µm. **(B)** Quantification of cyst diameter at 48 hours and **(C)** 96 hours following treatment. **(D)** Schematic for *in vivo* therapeutic evaluation. *Pkd1*^fl/fl^; Pax8-Cre mice were induced with doxycycline at P10 and intraperitoneally (I.P.) injected with CD-m*Pkd2* (1 mg/kg) at P13. Kidneys were harvested at P21 for analysis. Scale bar = 1 cm. **(E)** Gross kidney images across treatment groups. **(F)** Quantification of KW/BW from mice after different treatments. **(G)** Representative whole-kidney histological sections showing cystic morphology and **(H)** quantification of cystic index in PBS-treated, CD-m*Pkd2*–treated, and healthy mice. Scale bar = 2 mm. **(I)** Blood urea nitrogen (BUN) measurements at P21 across treatment groups. **(J)** Heatmap of RT–qPCR analysis showing expression of inflammatory and injury-associated genes (*Ccl2, Tnfa, Myc, Lcn1, Havcr1*) in kidneys from treated mice relative to PBS controls. Data are represented as mean ± SD. Statistical analysis was calculated with a one-way ANOVA with Tukey’s post hoc test. ns = not significant, * p < 0.05, ** p < 0.01, *** p < 0.001, **** p < 0.0001.

We next evaluated therapeutic efficacy in a rapidly progressive inducible *Pkd1*^fl/fl^;Pax8-Cre mouse model. Mice were induced at P10 and treated with CD-m*Pkd2* (1 mg/kg) at P13 (**Fig. 5D**). At P21, gross morphology revealed a reduction in renal enlargement compared to PBS-treated mice (**Fig. 5E**). Quantification demonstrated a 2.1-fold reduction in kidney weight–to–body weight (KW/BW) ratio relative to PBS controls (**Fig. 5F**). However, KW/BW ratios remained significantly elevated compared to uninduced healthy mice (p < 0.01), indicating partial but not complete normalization (**Fig. 5F**). Histological analysis corroborated these findings. CD-m*Pkd2* treatment reduced cystic burden and decreased cystic index by 48% relative to PBS-treated mice (**Fig. 5H**). Despite this improvement, cystic index remained higher than in healthy controls, consistent with partial structural rescue (**Fig. 5G, H**). Renal function was also improved with CD-m*Pkd2* administration. Serum blood urea nitrogen (BUN) levels were reduced by 46% in CD-m*Pkd2*–treated mice compared to PBS-treated controls, indicating functional benefit despite persistent structural abnormalities (**Fig. 5I**). Molecular analysis further supported partial disease attenuation. RT-qPCR demonstrated reduced expression of inflammatory and injury-associated genes, including *Ccl2*, Tnfa, Myc, *Lcn1*, and *Havcr1*, in CD-m*Pkd2*–treated kidneys relative to PBS controls (**Fig. 5J**). Although transcript levels remained elevated compared to healthy mice, expression patterns shifted toward a less cystogenic state (**Fig. 5J**). Collectively, these data demonstrate that PC2 supplementation via CD-m*Pkd2* significantly attenuates cyst growth and improves renal function in *Pkd1*-deficient models,

## Discussion

In this study, we developed a CD–targeted LNP (CD LNP) platform capable of delivering therapeutic mRNA to cyst-lining renal epithelial cells and demonstrated that PC2 replenishment can reverse established cystic kidney disease in *Pkd2*-deficient models and attenuate disease in *Pkd1*-deficient models (**Fig. 4**, **Fig. 5**). These findings address two central challenges in ADPKD therapy, efficient mRNA delivery to the kidney and restoration of polycystin function in cyst-forming epithelia. Efficient renal delivery has remained a major limitation of systemic LNP platforms, which typically exhibit dominant hepatic tropism^26^. By optimizing lipid composition and incorporating a CD-targeting peptide, we achieved enhanced renal accumulation and uptake within DBA-positive CD epithelia (**Fig. 3H–K**).

Although LNPs exceed the size threshold for glomerular filtration, their renal accumulation may be mediated through peritubular capillary access, as nanoparticles exceeding 15 nm have been shown to cross from the peritubular endothelium into renal tubular epithelium independently of glomerular filtration^32,33^. In ADPKD, disease-associated vascular remodeling, increased permeability, and skewed cell composition may further facilitate nanoparticle transport into the renal interstitium^34,35^. In addition, incorporation of cationic lipids such as DOTAP has been associated with enhanced kidney tropism, potentially through altered protein corona formation and endothelial interactions^24^. Once within the renal parenchyma, the CD-targeting peptide promotes uptake by CD epithelia, enabling delivery to cyst-lining cells. Although delivery was not exclusive to CD cells, enrichment within cyst-lining CD epithelia was sufficient to achieve biologically meaningful PC2 replenishment and therapeutic benefit. These results demonstrate that modest shifts in intrarenal targeting can translate into substantial disease modification in a genetically driven renal disorder.

Notably, repeated administration of CD-m*Pkd2* at 1 mg/kg induced regression of established disease, with kidneys exhibiting architectural, cell composition, and transcriptional features approaching those of healthy controls (**Fig. 4B, F, G**). Consistent with prior studies demonstrating that re-expression of *Pkd2* can reverse cyst burden^19^, our findings show that restoration of renal PC2 with CD-m*Pkd2* similarly induces regression of established disease. These results support the concept that polycystin deficiency remains a reversible driver of cyst maintenance, not solely initiation, and highlight PC2 restoration using CD-m*Pkd2* as a potentially translatable therapeutic strategy.

Replenishment of renal PC2 reduced expression of inflammatory mediators (*Ccl2, Tnfa*), fibrotic markers (*Acta2*), and injury-associated transcripts (*Havcr1*, *Lcn1*) (**Fig. 4H**), coinciding with reduced macrophage infiltration and myofibroblast accumulation (**Fig. 4G**). These findings suggest that correcting the primary defect can remodel the pericystic niche, which is increasingly recognized as a contributor to cyst progression^36^. Moreover, repeated dosing was well tolerated, with no detectable elevations in AST or ALT and no histological abnormalities in liver, lung, spleen, or heart (**Fig. S4**). Although these findings support a favorable short-term safety profile, longer-term studies will be necessary to evaluate durability of therapeutic response, potential immunogenicity, and cumulative toxicity. The requirement for repeated administration also raises important considerations regarding dosing frequency and durability of PC2 expression.

In *Pkd1*-deficient kidneys, miR-17 suppresses endogenous *PKD2* translation via 3’UTR binding sites, limiting endogenous PC2 compensation^20,21,37^. Exogenous m*Pkd2* lacks these binding sites and thus escapes this repression. Importantly, Lakhia et al. demonstrated that increasing PC2 levels through miR-17 inhibition attenuates cyst progression in *Pkd1*-deficient mice, establishing that PC2 supplementation can mitigate *PKD1*-deficient disease^20^. Our findings extend this work by showing that transient, CD-m*Pkd2*–mediated restoration of PC2 reduces cyst burden and improves renal function in a rapid inducible *Pkd1* model (**Fig. 5E–J**). This effect may reflect, in part, the ability of PC2 to partially compensate for PC1 loss through restoration of calcium signaling and modulation of downstream cAMP-dependent pathways, as well as residual formation or stabilization of polycystin complexes^20,38^. Despite attenuating disease progression in *Pkd1-*deficient mice, CD-m*Pkd2* mice still presented with elevated KW/BW, cyst burden, and BUN compared to healthy mice (**Fig. 5E-J**). This finding suggests that PC1 deficiency imposes additional constraints that limit the extent of rescue achievable by PC2 supplementation alone. The degree of PC2 restoration required for optimal therapeutic benefit in *Pkd1-*driven disease remains to be defined and may necessitate higher or sustained levels of PC2 expression.

Despite these promising results, several limitations should be acknowledged. First, although CD LNPs enhanced uptake in CD cells, delivery remained distributed across multiple renal cell populations. While this did not preclude therapeutic efficacy, further refinement may improve cell-type specificity, reduce off-target exposure, and enhance dose efficiency. In addition, CD LNPs exhibited hepatic uptake, suggesting that optimization of lipid composition through *in vivo* distribution studies may further improve organ specificity. Although no evidence of hepatotoxicity was observed and liver accumulation did not limit therapeutic efficacy, increased kidney specificity may further reduce off-target exposure and lower the dose required for therapeutic benefit. Second, the durability of PC2 restoration was not evaluated. Given the transient nature of mRNA-mediated protein expression, defining the duration of PC2 restoration will be critical for optimizing dosing frequency to maximize therapeutic efficacy while minimizing systemic exposure. In contrast to viral gene therapies that enable sustained expression, our approach will require repeated administration to maintain therapeutic PC2 levels. Third, although CD-m*Pkd2*–mediated PC2 supplementation attenuated disease in *Pkd1*-deficient models, rescue was incomplete, indicating that higher or sustained levels of PC2 may be required for maximal therapeutic benefit in *PKD1*-deficient disease. Moreover, our studies utilize the well-characterized *Pkd2*^fl/fl^; Pax8-Cre and Pkd1^fl/fl^; Pax8-Cre mouse models, which recapitulate cystogenesis driven by near complete gene inactivation. While these models are widely used for preclinical therapeutic evaluation in ADPKD, some patients harbor hypomorphic alleles and retain partial *PKD1* or *PKD2* function^39,40^. Accordingly, future studies will prioritize evaluating therapeutic efficacy in alternative ADPKD models that more closely reflect the human genetic landscape, including *Pkd2*^WS25/-^ and Pkd1^RC/RC^ strains^41–43^.

Overall, our findings establish that kidney-directed *Pkd2* mRNA replacement can modify, and in *Pkd2*-driven disease, reverse, established ADPKD pathology. Unlike current therapies that act downstream of polycystin loss, this strategy directly addresses the underlying molecular defect. More broadly, these results provide a translational framework for mRNA delivery to renal epithelia and support the feasibility of gene-based therapy for monogenic kidney disorders.

## Methods

### Materials

D-lin-MC3-DMA (MC3), ALC-0315 (ALC), DSPE-PEG(2000)-maleimide, cholesterol, DOTAP (1,2-dioleoyl-3-trimethylammonium-propane), and DSPE-PEG(2000)-methoxy (1,2-distearoyl-sn-glycero-3-phosphoethanolamine-N-[methoxy(polyethylene glycol)-2000] were purchased from Avanti Polar Lipids (Alabaster, AL, USA). Lipids were dissolved in ethanol prior to nanoparticle formulation. The CD–targeting peptide (CKDSPKSSKSIRFIPVST) was synthesized by BioMatik (Ontario, Canada). mCherry mRNA, luciferase mRNA (mLuc), and full-length mouse and human *Pkd2* mRNA (m*Pkd2*) were obtained from TriLink BioTechnologies (San Diego, CA, USA) and included a 5′ cap structure and a 300-adenosine poly(A) tail. All other reagents were obtained from Sigma-Aldrich unless otherwise specified.

### Lipid Nanoparticle Synthesis

LNPs were synthesized using hand mixing as previously described^44^. Lipids dissolved in ethanol were combined with mRNA diluted in citrate buffer (50 mM, pH 4.0) at a defined ionizable lipid-to-mRNA weight ratio. The ethanol and citrate buffer phases were mixed at a 3:1 (v/v) ratio. For CD LNPs, prior to synthesis, the cysteine-containing CD-targeting peptide was conjugated to DSPE-PEG(2000)-maleimide as previously described^29,45–47^. Briefly, DSPE-PEG(2000)-maleimide was mixed with the CD-targeting peptide overnight at pH 7.4 and then purified through high-performance liquid chromatography (HPLC, Prominence, Shimadzu, Columbia, MD, USA) on a Jupiter C4 reverse-phased column (Phenomenex, Torrance, CA, USA) at 55 °C using water and acetonitrile supplemented with 0.1% formic acid as the mobile phases. The molecular weight of the eluted peptide-lipid amphiphile was characterized using matrix-assisted laser desorption/ionization time-of-flight mass spectroscopy (MALDI-TOF-MS, Bruker, MA, USA). The resulting DSPE-PEG(2000)-CD-targeting peptide was incorporated into LNP formulations in place of DSPE-PEG(2000)-methoxy at specified molar percentages. Following mixing, LNPs were dialyzed against sterile PBS to remove ethanol, citrate buffer, and unincorporated lipids. Prior to use, LNPs were sterile filtered through 0.22 μm filters.

### LNP Characterization

Hydrodynamic diameter, zeta potential, and polydispersity index were measured using dynamic light scattering on a Zetasizer Ultra (Malvern Panalytical, Malvern, UK). Encapsulation efficiency of mRNA was determined using a RiboGreen RNA assay (Thermo Fisher Scientific) with and without treatment with Triton X-100 to disrupt nanoparticles. For gel shift assays, mRNA LNPs were mixed with loading buffer and run on a 2% agarose gel at 60V for 30 minutes to confirm mRNA encapsulation. For TEM, LNPs (1 µg/mL mRNA) were placed onto copper 200-mesh carbon grids for 10 minutes and then stained with 2 % (w/v) uranyl acetate for 10 min. Grids were left overnight to dry and imaged on a Talos TEM at 80 kV (Thermo Fisher Scientific, Waltham, MA, USA).

### Cell Culture

Mouse principal cortical collecting duct (CD) cells and proximal tubule (PT) epithelial cells, inner medullary collecting duct (IMCD) cells derived from inducible *Pkd2* knockout mice, and *Pkd1* knockout IMCD cells were routinely cultured in RenaLife™ epithelial medium complete kit (Lifeline, Frederick, MD, USA). Cells were incubated at 37 °C in a humidified incubator containing 5% CO₂. IMCD cells derived from inducible *Pkd2* knockout mice were provided by the PKD Research Resource Consortium (PKD RRC, Baltimore, MD, USA). *Pkd1* knockout IMCD cells were previously characterized and provided by Stefan Somlo’s lab at Yale University^48^.

### In vitro Transfection

To evaluate transfection efficiency, cells were treated with LNPs encapsulating mCherry mRNA at concentrations ranging from 125–1000 ng/mL. After 24 hours, mCherry expression was quantified using fluorescence microscopy and flow cytometry. Cellular metabolic activity following nanoparticle treatment was assessed using an MTS assay according to the manufacturer’s instructions. For competition assays, cells were preincubated with free CD-targeting peptide for 30 minutes at 50 µM prior to treatment with DiR-labeled LNPs.

### Immunofluorescence

Cells were fixed with 10% formalin and permeabilized with 0.1% Triton X-100. Samples were blocked with 2% bovine serum albumin and incubated with primary antibodies against polycystin-2 (PC2), DBA lectin for CD cells, or LTL for proximal tubules. After washing, samples were incubated with fluorophore-conjugated secondary antibodies and counterstained with DAPI. Images were acquired using a confocal fluorescence microscope and analyzed using ImageJ software.

### RT-qPCR

Total RNA was extracted from cultured cells or kidney tissue using TRIzol reagent. cDNA was synthesized using a reverse transcription kit (Qiagen, Germantown, MD, USA). Quantitative PCR was performed using SYBR Green Master Mix on a QuantStudio real-time PCR system. Gene expression levels were normalized to the *Gapdh* gene and analyzed using the ΔΔCt method. Target genes included *Ccl2, Tnfa, Ano1, Myc, Lcn1, Havcr1, and Acta2*.

### 3D Cystogenesis Assays

For *in vitro* cyst formation assays, IMCD cells were embedded in Matrigel and cultured in cystogenic media supplemented with forskolin (10 µM, ThermoFisher, Cat. No: J63292.MF), as previously described^49–51^. Cells were treated with LNP formulations delivering mRNA at defined concentrations. Cyst diameter was measured at 48 and 96 hours after treatment using brightfield microscopy. Quantification was performed using ImageJ.

### Animal Studies

All animal procedures were approved by the Institutional Animal Care and Use Committee (IACUC) at the University of Southern California. Inducible *Pkd2*^fl/fl^;Pax8-Cre and *Pkd1*^fl/fl^;Pax8-Cre mouse models were used to evaluate therapeutic efficacy. Gene deletion was induced via doxycycline administration at defined developmental time points. For biodistribution studies, *Pkd2*^fl/fl^;Pax8-Cre mice received a single intravenous injection of DiR-labeled nanoparticles on P125. Then, on P126, organs were harvested and imaged using an IVIS Spectrum imaging system.

### Therapeutic Evaluation Studies

For reversal studies, *Pkd2*^fl/fl^;Pax8-Cre mice were induced with doxycycline from P28-31. Starting on P99, mice received repeated CD-m*Pkd2* administration (tail vein) every three days for four weeks. Kidney weight-to-body weight (KW/BW) ratios, cystic index, and histological markers were evaluated. To evaluate the therapeutic efficacy in a *Pkd1*-deficient model, *Pkd1*^fl/fl^;Pax8-Cre mice were induced with doxycycline at P10 and then received CD-m*Pkd2* at 1 mg/kg at P13. Mice were euthanized at P21 for tissue analysis^33,51^.

### Histology and Immunohistochemistry

Organs were embedded in optimal cutting temperature (OCT) compound, frozen in 2-methylbutane cooled with liquid nitrogen, and cryosectioned at 10 μm thickness using a Leica CM3050S cryostat (Leica, Wetzlar, Germany). Sections were then stained with hematoxylin and eosin (H&E) or Sirius red, according to the manufacturer’s instructions. Images were captured using a Leica DMi8 microscope (Leica, Wetzlar, Germany). For immunohistochemistry, fresh sections were fixed with 10% formalin for 10 minutes, permeabilized with 0.1% triton-X100 for 30 minutes, and blocked with 2% BSA for 30 minutes. Following several washes, sections were incubated with primary antibodies against macrophages (F4/80, 1:100) and myofibroblasts (α-SMA, 1:200) overnight at 4°C. The following day, samples were incubated with a fluorophore conjugated secondary antibody (1:1000) and DAPI (1:1000) for 1 hour at room temperature. Images were captured using a Leica DMi8 microscope (Leica, Wetzlar, Germany).

### Flow Cytometry

Kidneys were dissociated into single-cell suspensions as previously described^52^. Briefly, kidney tissue was minced and incubated with collagenase (1 mg/mL) and dispase (1 mg/mL) for 1 hour at 37°C. Digest was passed through a 40 µm cell strainer, pelleted at 300 g for 5 min, and red blood cells (RBC) were lysed using ammonium chloride-based RBC lysis buffer. The remaining cells were pelleted at 300 g for 5 min and incubated with DAPI (to exclude dead cells) and antibodies or lectins to identify specific renal cell populations (see table 1). Data were acquired using a CytoFLEX cytometer (Beckman Coulter, Brea, CA, USA) and analyzed using FlowJo software. Single-cell suspensions were gated on singlets and DAPI-negative populations before subset identification using lectin- and antibody-based markers. LNP-positive cells were defined by DiR fluorescence above the signal threshold established in unlabeled controls, and compensation was performed using single-stained controls.

### Serum Markers and Toxicity Assessment

Blood samples were collected at experimental endpoints. Serum levels of alanine aminotransferase (ALT), aspartate aminotransferase (AST), and blood urea nitrogen (BUN) were measured according to the manufacturer’s instructions to evaluate liver and kidney function.

### Statistical Analysis

All experiments were performed with at least two biological replicates unless otherwise specified. Data are presented as mean ± standard deviation. Statistical analyses were performed using GraphPad Prism software. Comparisons between two groups were performed using unpaired two-tailed Student’s t-tests, while multiple group comparisons were conducted using one-way or two-way ANOVA with appropriate post hoc tests. Differences were considered statistically significant at p < 0.05.

### Antibodies

| Name | Dilution | Catalog Number |
| --- | --- | --- |
| Dolichos Biflorus Agglutinin (DBA), Fluorescein | 1:200 | Vector Laboratories (FL-1031-2) |
| Lotus Tetragonolobus Lectin (LTL), Fluorescein | 1:200 | Vector Laboratories (FL-1321-2) |
| Polycystin 2 Polyclonal antibody | 1:100 | Proteintech (19126-1-AP) |
| Alpha-Smooth Muscle Actin Monoclonal Antibody (1A4) | 1:200 | ThermoFisher (14-9760-82) |
| F4/80 Monoclonal Antibody | 1:100 | ThermoFisher (MA5-16363) |
| PEG Monoclonal Antibody | 1:100 | GenScript (A01795) |
| Goat anti-Rabbit IgG (H+L) Cross-Adsorbed Secondary Antibody, Alexa Fluor™ 594 | 1:1000 | ThermoFisher (A-11012) |
| CD140b (PDGFRB) Monoclonal Antibody (APB5), PE | 1:100 | ThermoFisher (12-1402-81) |
| CD45 Monoclonal Antibody (30-F11), FITC | 1:100 | ThermoFisher (11-0451-82) |
| F4/80 Monoclonal Antibody (BM8), APC | 1:100 | ThermoFisher (17-4801-82) |
| CD31 (PECAM-1) Monoclonal Antibody (390), FITC | 1:200 | ThermoFisher (11-0311-82) |
| Wheat Germ Agglutinin (WGA)-Alexa Fluor 488 | 1:100 | ThermoFisher (W11261) |

---

**Figure S1:**
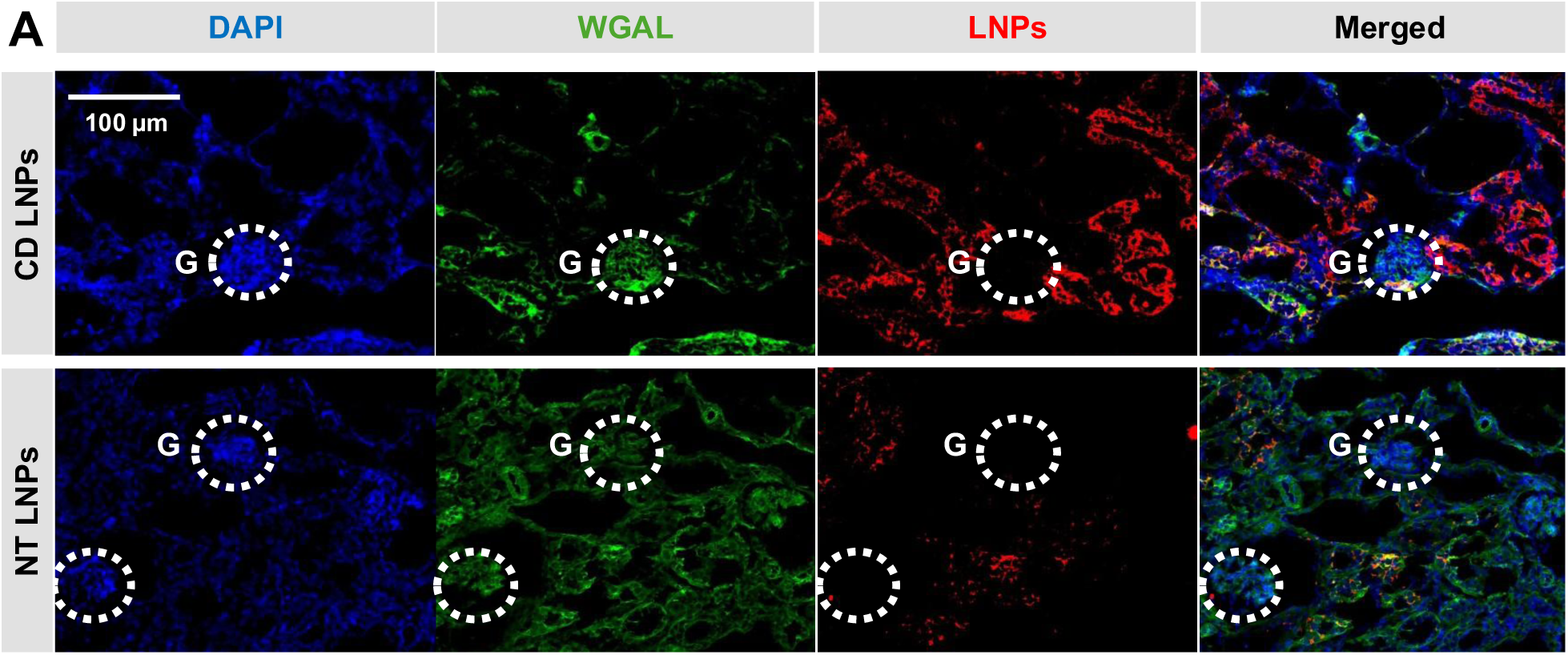
Distribution of CD LNPs in *Pkd2*-deficient kidneys. (**A**) Representative immunofluorescence images of kidney sections following administration of DiR-labeled CD LNPs or NT LNPs. Nuclei are stained with DAPI (blue), nephrons with WGA lectin (green), and LNPs are shown in red. Merged images highlight colocalization patterns. Glomeruli (G) are indicated by dashed circles. Scale bar = 100 μm.

**Figure S2:**
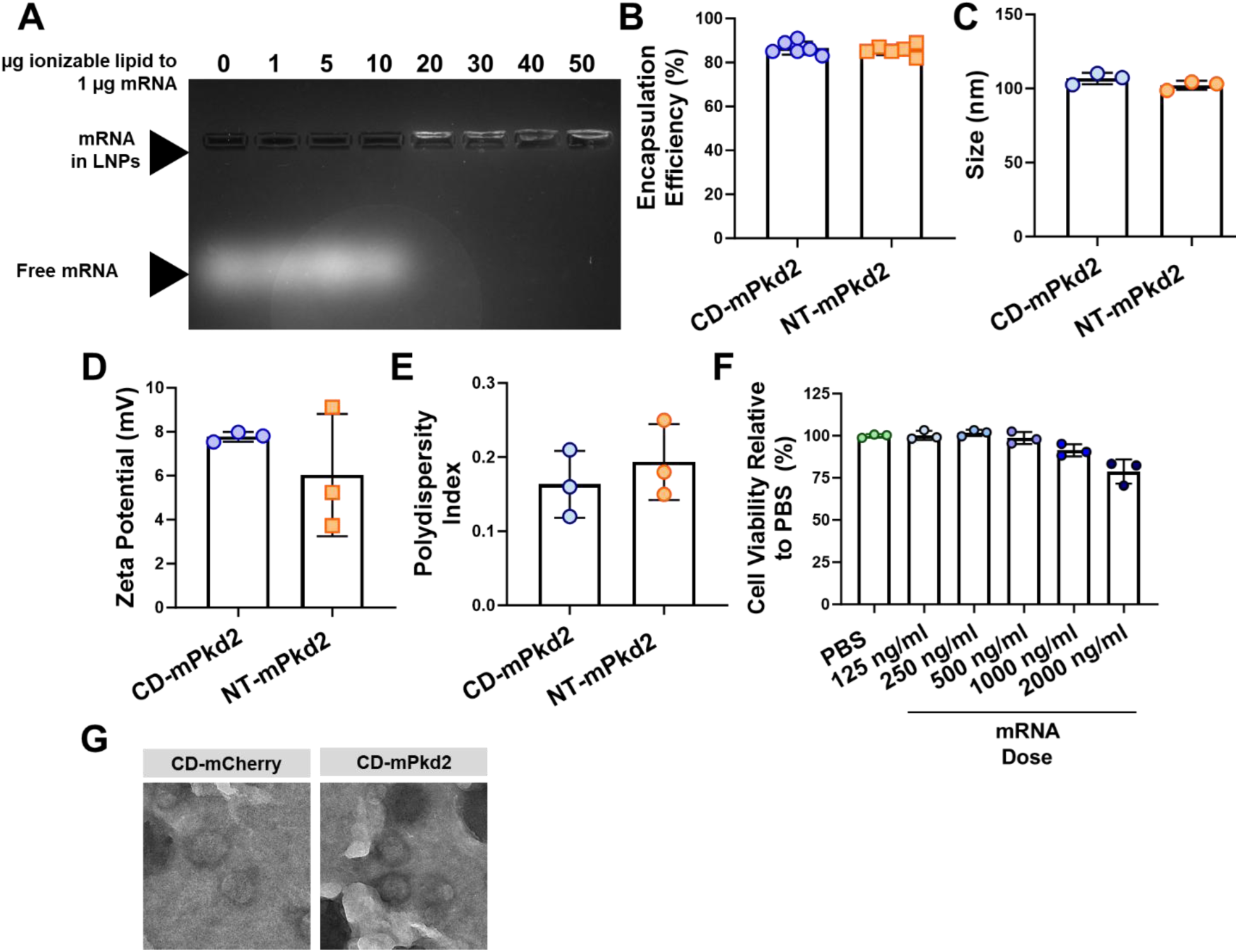
Characterization of CD-m*Pkd2* LNPs. (**A**) Agarose gel electrophoresis of mRNA cargo in LNP formulations made at different ionizable lipid to mRNA weight ratios. (**B**) Encapsulation efficiency (%), **(C)** hydrodynamic diameter (size (nm)), **(D)** zeta potential (mV), and **(E)** polydispersity index (PDI) values of CD- and NT-m*Pkd2* LNPs. (**F**) Cell viability in IMCD cells after 48-hour incubation with various doses of CD-m*Pkd2*. (**G**) Transmission electron microscopy images of CD LNPs encapsulating control mCherry mRNA (CD-mCherry) or therapeutic m*Pkd2* mRNA (CD-m*Pkd2*), confirming spherical morphology and comparable nanostructure between formulations. N = 3. Data are presented as mean ± SD.

**Figure S3:**
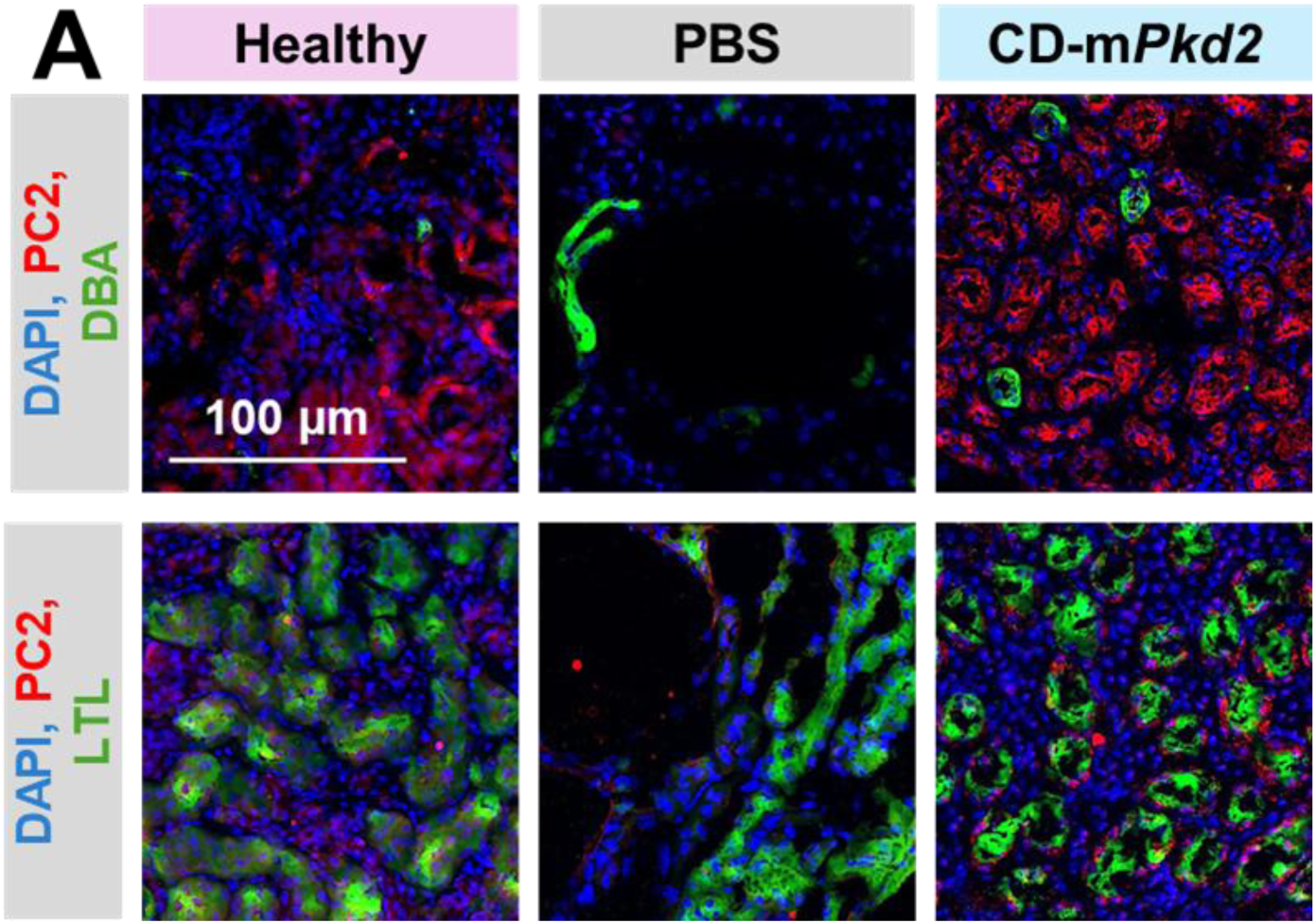
Polycystin-2 expression in *Pkd2*-deficient kidneys after repeated administration of CD-m*Pkd2*. (**A**) *Pkd2*^fl/fl^; Pax8-Cre mice were induced with doxycycline from P28-31. Mice received intravenous (I.V.) injections of CD-m*Pkd2* at 1.0 mg/kg or PBS beginning at P99 every three days until P128. Kidney sections stained for PC2 (red), DBA (top row, green), LTL (bottom row, green), and DAPI (nuclei, blue) show that kidneys treated with 1 mg/kg CD-m*Pkd2* had elevated PC2 signal compared to PBS-treated mice. N = 4. Scale bar = 100 μm.

**Figure S4:**
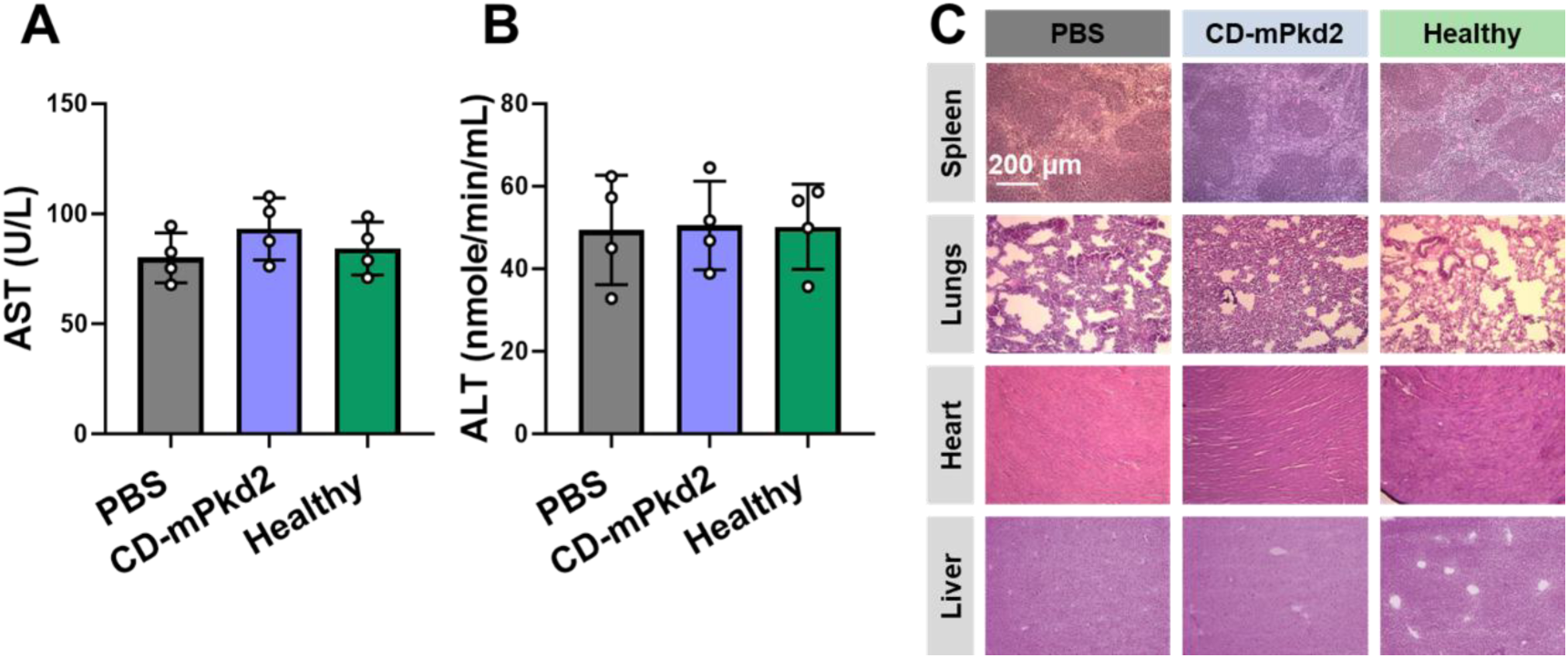
Liver enzymes and extrarenal organ morphology after repeated administration of CD-m*Pkd2*. (**A-B**) Serum analysis of liver function (AST and ALT) markers following administration of CD-*mPkd2* compared to PBS and healthy controls, showing no significant changes across groups. **(C)** Representative H&E staining of major organs (spleen, lungs, heart, and liver) collected after treatment with PBS CD-m*Pkd2* (1 mg/kg) LNPs, alongside healthy controls. No overt histopathological abnormalities were observed across treatment groups. Scale bar = 200 μm. Data are presented as mean ± SD.

